# Molecular Dynamics-Guided All-Atom Reconstruction of Cryo-ET Maps Reveals Mechanisms of Histone Tail-Mediated Chromatin Compaction

**DOI:** 10.1101/2025.11.05.686627

**Authors:** Shuxiang Li, Maria J. Aristizabal, Sergei A. Grigoryev, Anna R. Panchenko

## Abstract

Dynamics and physical state of chromatin are crucial in regulating gene expression, DNA replication, and repair. Intrinsically disordered histone tails were previously recognized as key modulators of chromatin states. However, detailed atomistic mechanisms by which histone tail dynamics are associated with chromatin compaction and higher-order chromatin organization remain poorly understood. In this work, we combine extensive all-atom molecular dynamics simulations of tri-nucleosomes with cryo-electron tomography (Cryo-ET) of native nucleosome arrays to investigate histone tail-mediated chromatin folding. Our approach offers distinct advantages as it elucidates realistic inter- and intra-nucleosomal interactions and DNA conformations derived from physics-based MD simulations, enabling a more accurate and physically grounded interpretation of Cryo-ET data. The results reveal that histone tails promote chromatin compaction and constrain tri-nucleosome unfolding via three major patterns: through histone-DNA interactions, histone H2A-H4 and H3-H4 tail-tail interactions. Notably, the distributions of MD-generated structural parameters of tri-nucleosomes with truncated histone tails were found to be in strong agreement with those of experimental open chromatin arrays with widely spaced nucleosomes, whereas the system with histone tails resembled more condensed chromatin. These findings provide mechanistic insights into how histone tails may mediate chromatin folding at the atomistic scale and underscore the dual role of histone tails in both structural compaction and inter-nucleosomal communication.

## Introduction

Chromatin, the complex of DNA and histone proteins, plays a central role in regulating cellular processes such as gene expression, DNA replication, transcription, recombination, and DNA repair. The dynamic folding and unfolding of chromatin are essential for maintaining its functional state, with histone N- and C-terminal tails playing a central role in these processes ^1,2^. Histone tails, which are intrinsically disordered protein regions, participate in various interactions that facilitate chromatin compaction, the higher-order organization of chromatin fiber and stability ^3–5^.

Understanding the physical principles governing chromatin folding remains a key challenge in chromatin biology ^6–8^. A number of experimental techniques, including Cryo-electron microscopy (Cryo-EM), Cryo-electron tomography (Cryo-ET), X-ray scattering, super-resolution fluorescence microscopy, NMR and chromosome conformation capture methods, have been employed to explore the structural organization of chromatin both *in vitro* and *in vivo* ^9–16^. While these methods have provided crucial insights into chromatin architecture, achieving atomic-level resolution remains difficult, particularly for histone tails in condensed chromatin regions. This limitation arises from the intrinsic flexibility of chromatin and the transient, heterogeneous nature of histone tail interactions with both nucleosomal and linker DNA ^17–19^. An early structural model of chromatin posited that histone tails may mediate inter-nucleosomal interactions necessary for forming compact fibers ^9,20^. In particular, interactions between the H4 tail and the acidic patch of H2A-H2B dimers have been extensively studied ^21^, but a full mechanistic understanding of how different histone tails contribute to chromatin compaction, especially through DNA-mediated or multi-nucleosome interactions, remains incomplete ^22^.

Another major challenge involves reconstructing realistic 3D chromatin models from Cryo-EM or ET maps. Traditionally, this has been achieved by rigidly fitting mono-nucleosome crystal structures into high-density volumes of Cryo-ET or EM data ^9,11,12^. However, such approaches suffer from significant limitations: they rely on arbitrary decisions about nucleosome chain direction, fail to capture the spatial coherence of nucleosome arrays, and often connect nucleosomes using straight DNA linkers that ignore the geometric and torsional constraints of DNA. These issues are exacerbated in highly condensed chromatin regions, where ambiguous fits and poor DNA resolution complicate the generation of reliable models.

To overcome these limitations and provide mechanistic insight into chromatin folding, we employed all-atom molecular dynamics (MD) simulations of tri-nucleosomes ̶ the minimal chromatin unit capable of capturing inter-nucleosomal geometry, linker DNA conformations, and histone tail interactions. We demonstrate that these tri-nucleosome models can be used to allow for the near-atomic interpretation of Cryo-ET density maps. Namely, these models specify the relative orientation of consecutive nucleosomes while simultaneously and inherently incorporating DNA and protein geometric and torsional constraints, even in partially resolved maps. This tri-nucleosome-based fitting framework provides a conceptually novel and practically superior strategy for building biologically meaningful chromatin models. Additionally, we present a systematic analysis of tri-nucleosome dynamics in systems with and without histone tails and show that histone tails are critical for driving chromatin compaction, stabilizing folded conformations, and mediating inter-nucleosomal interactions. Importantly, we find that the tri-nucleosome conformations generated in our simulations reproduce key stereological features of experimental native chromatin. Moreover, we reveal previously uncharacterized modes of histone tail interaction that provide new mechanistic insights into how chromatin structure is regulated.

## Results

### Chromatin Modeling Based on Tri-nucleosomes Simulations Offers Distinct Advantages

Previous efforts to model chromatin and nucleosome arrays have predominantly relied on fitting a nucleosome core particle X-ray crystal structure into the high-density regions of Cryo-ET or tomographic density maps ^9,11,12,23,24^. Mono-nucleosome based fitting has certain advantages, such as a more accurate local placement since the local measure cross-correlation coefficient is maximized (see Methods). This is reflected in relatively high values of the local goodness-of-fit measures (Figures S4c, S5c). However, mono-nucleosome based fitting strategies have several inherent limitations. One major drawback is the arbitrary selection of the nucleosome chain path, which often leads to ambiguous or inconsistent definitions of consecutive nucleosome arrangement (Figure 1a, see Figures S1 for strong contrast with black background and volume rendering of Cryo-ET density maps). Additionally, straight-line DNA segments are typically used to connect nucleosomes that ignore the torsional mechanics and geometric constraints of DNA, such as twist, rise, and bending, that are essential for maintaining the structural integrity and topological regulation of chromatin ^25^. Another major limitation of mono-nucleosome fitting occurs in the condensed chromatin regions within the Cryo-ET map, where multiple mono-nucleosomes can be erroneously fitted into the same high electron density volume area (Figure 1c). This necessitates manual intervention and subjective decision-making to determine reasonable nucleosome configurations. Moreover, due to the varying and crowded density volumes of Cryo-ET maps, tracing all linker DNA regions becomes especially challenging in condensed chromatin.

**Figure 1.**
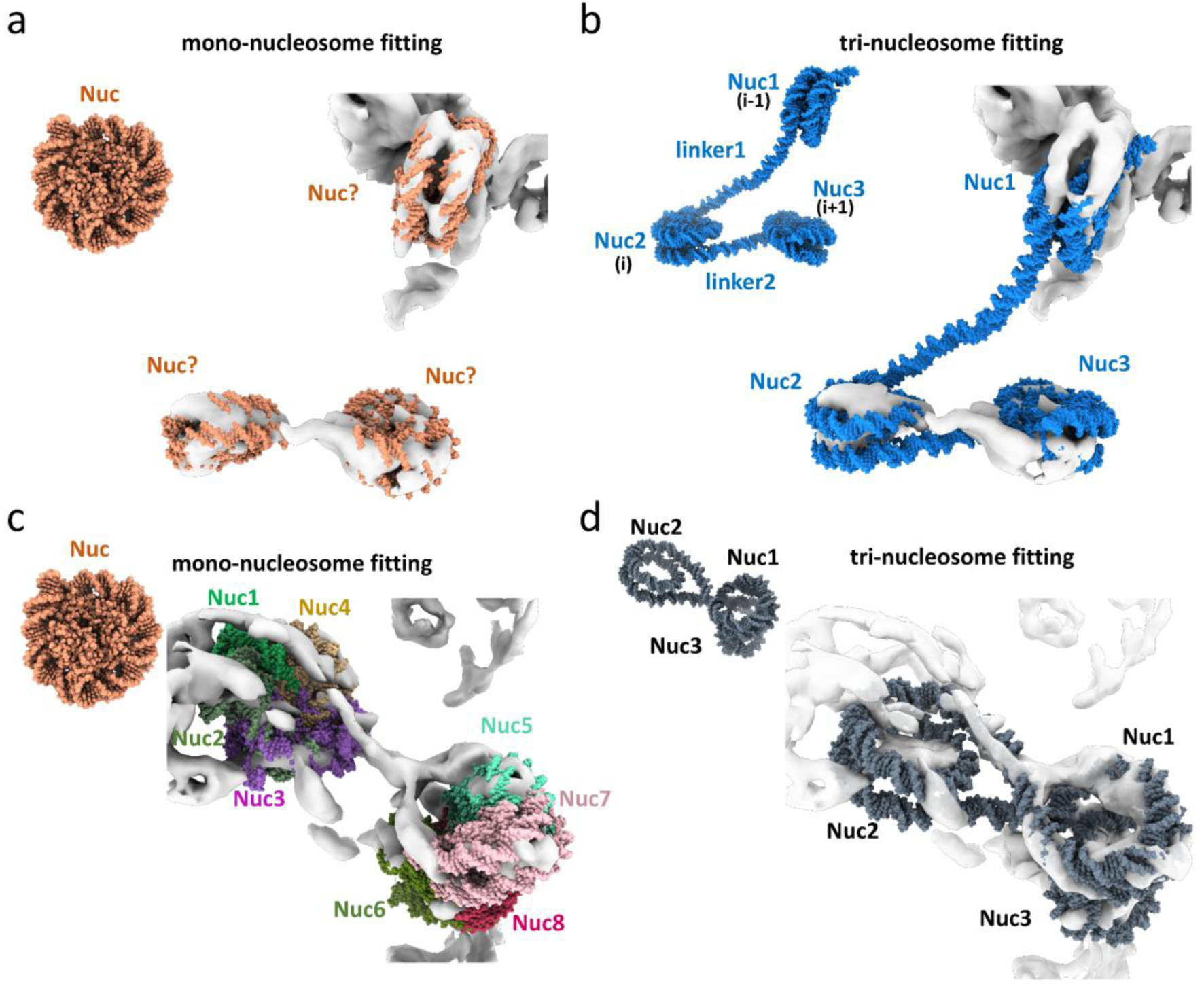
Comparative analysis of fitting of mono-nucleosome and tri-nucleosome models into cryo-ET density maps of human chromatin. **a** Fitting of a mono-nucleosome structure (PDB: 1KX5) into high-density regions of the cryo-ET map of an open chromatin region (from panel 10 of Supplementary Figures S10 with the crosslinked chromatin array vitrified at 0 mM Mg^2+^). **b** Fitting of a representative tri-nucleosome conformation from NUC_notail_ MD simulations into the same cryo-ET map as shown in (a). **c** Fitting of the mono-nucleosome structure into the cryo-ET map of a condensed chromatin region (from panel 1 of Supplementary Figures S3 with the chromatin array vitrified at 0.75 mM Mg^2+^). **d** Fitting of a representative tri-nucleosome conformation from NUC_tail_ MD simulations into the same cryo-ET map as shown in (c).

Our approach utilizes tri-nucleosome conformations derived from MD simulations to fit into the Cryo-ET maps. These conformations inherently define the relative spatial organization and order of three consecutive nucleosomes (Nuc1, Nuc2, Nuc3), providing information about nucleosome arrangement and a more meaningful representation of local regions of chromatin and nucleosome clutches (Figure 1b). Importantly, since these tri-nucleosome conformations are sampled from physically plausible MD trajectories, the associated linker DNA segments naturally embody torsional strain and geometric constraints, offering a more accurate structural conformation and interpretation for modeling of nucleosome arrays. Moreover, when tri-nucleosome core structures are aligned with appropriate high-density regions in the map, the linker DNA segments connecting the three nucleosomes can be positioned in alignment with the corresponding DNA density, even when the linker DNA densities are partially missing or poorly resolved in the Cryo-ET map (Figure 1d). As such, tri-nucleosome fitting can improve the model’s physical plausibility and provide insight into the geometry and connectivity of nucleosome arrays within chromatin.

### Tri-nucleosome Simulations Provide Near-Atomistic Models for Experimental Chromatin Maps

First, we assessed the goodness of fit of Cryo-ET maps using the MD-generated tri-nucleosome conformations. Cryo-ET maps were collected under two experimental conditions: 0 mM Mg²⁺ (open chromatin architecture) and 0.75 mM Mg²⁺ (condensed chromatin architecture), the latter approximating the physiological concentration of free magnesium in a cellular nucleus ^12,26^ (Figures S2 and S3). We found that fitting of the MD conformations of the tri-nucleosome system without tails (NUC_notail_) into the nucleosome array, captured at 0 mM Mg^2+^, yielded relatively high values for goodness-of-fit measures, like correlation coefficients and “inside-the-contour”, calculated for the whole tri-nucleosome system (Figures S4a, S5a). Using the MD-generated tri-nucleosome with tails’ conformations (NUC_tail_) yielded lower correlation coefficients and reduced inside-the-contour values compared to a system without tails (Figure S4b, S5b). These results suggest that the conformations of tri-nucleosome systems with histone tails are not compatible with the unfolded open chromatin architectures, whereas the tri-nucleosomes without tails conformations closely resemble such unfolded conformations. Conversely, we found that the NUC_tail_ conformations better matched the condensed chromatin structures when compared to the NUC_notail_ conformations (Figure S6-7).

To further assess the relevance and accuracy of the tri-nucleosome conformations, we calculated and compared the following structural characteristics: Euclidean distance between the disjointed nucleosomes (Nuc1–Nuc3, corresponding to N-distance in ^12^) and distance between the consecutive (Nuc1–Nuc2 or Nuc2–Nuc3, corresponding to D-distance) nucleosomes. In both cases, the experimental data were taken from the previous study ^12^ and the results of MD simulations were directly compared to experiments without any fitting involved. The distributions of MD-generated parameters were found to be in strong agreement with those of experimental chromatin arrays at 0 mM Mg²⁺, open chromatin with widely spaced nucleosomes. The disjointed nucleosome distance distribution had a median value of around 200 Å, whereas the consecutive nucleosome distribution showed a slightly larger median, ∼218 Å (Figure 2a-b, Figure S8-9).

**Figure 2.**
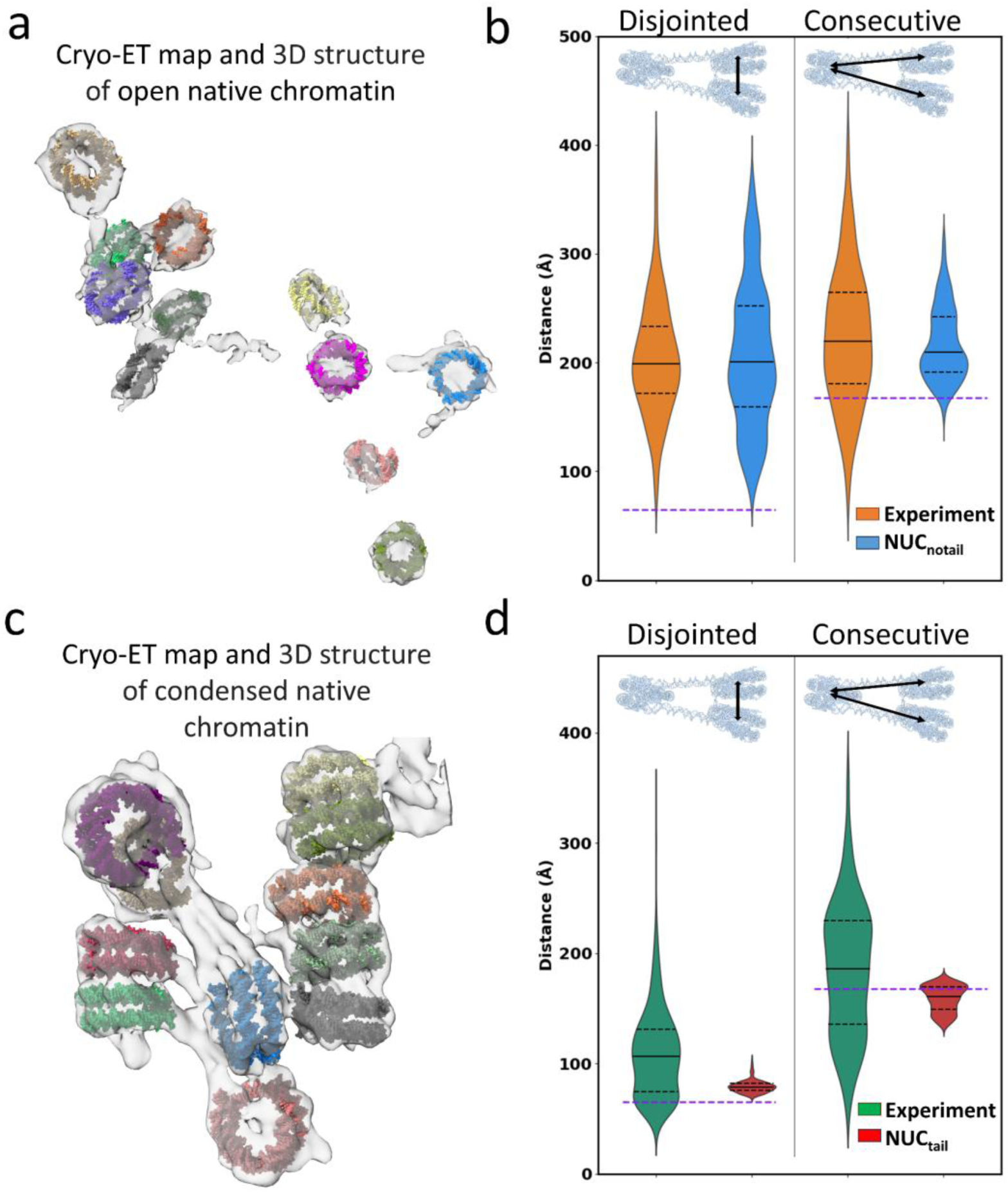
Tri-nucleosome MD simulations recapitulate chromatin conformations observed in Cryo-ET reconstructions. **a** 3D reconstruction of nucleosome arrays from cryo-ET density maps (gray) of open chromatin at 0 mM Mg²⁺ from ref ^10^. **b** Distribution of disjointed and consecutive nucleosome center-to-center distances measured from experimental chromatin (orange) and from NUC_notail_ simulations (blue) from all three runs. The purple dashed line shows the distance in the initial NUC_notail_ system (PDB: 6L4A). **c** 3D reconstruction of nucleosome arrays from cryo-ET density maps (gray) of condensed chromatin at 0.75 mM Mg²⁺ from ref ^10^. **d** Distribution of disjointed and consecutive center-to-center nucleosome distances from experimental chromatin (green) and from NUC_tail_ simulations (red). The purple dashed line shows the distance in the initial NUC_tail_ system.

Nucleosomal distances derived from experiments were much smaller in condensed chromatin arrays compared to open ones (without magnesium), reflecting the Mg²⁺-induced chromatin compaction (Figures 2c-d) ^12,27^. A distance reduction was also observed in the NUC_tail_ system, where chromatin unfolding was significantly suppressed due to the histone tail-mediated interactions, highlighting the role of histone tails in compacting chromatin arrays. It is noteworthy that the distance distributions of the NUC_tail_ system were narrower than experimental ones (Figure 2d). In addition, the inter-nucleosomal distances of the crosslinked chromatin displayed a slight reduction in disjointed nucleosomal distances while maintaining similar consecutive nucleosome distances, suggesting a modest compaction of Nuc1–Nuc3 nucleosomes (Figure S10-12).

### Structural and Dynamic Determinants of the Model’s Predictive Power

In the previous section, we showed that the tri-nucleosome system without tails serves as a suitable model for open chromatin. Here, we analyze the structural and dynamic features that underlie the model’s predictive power. In the NUC_notail_ system during MD simulations, pronounced unfolding was observed on one or both flanking nucleosomes with respect to Nuc2, which was used as a reference for aligning MD trajectories/frames (Figure S13). The process of unfolding involved the detachment and unwrapping of DNA ends from histone octamers, leading to an increase in the effective linker lengths. Indeed, even though the original linkers have 22 bp, MD trajectories showed a wide range of the effective linker lengths (Figure 2b). Indeed, if we subtract the nucleosomal diameter of 97 Å from the median value of center-to-center distance (218 Å) and divide by 3.4 Å (each base pair of DNA spans 3.4 Å), we will get an estimate of the effective linker lengths ranging from 13 bp to 68 bp with a median of 36 bp. This, in part, explains the consistency between the experimental and modeled consecutive nucleosomal distances discussed in the previous section, as in vivo chromatin fibers utilize linkers of varying lengths.

To quantify the DNA unwrapping at the individual nucleosome level, we calculated the radius of gyration (Rg) distributions for each nucleosome. In the NUC_notail_ system, both Nuc1 and Nuc3 exhibited bimodal Rg distribution with the first peak pointing to the DNA breathing and the second peak pointing to the extensive DNA unwrapping at the nucleosome entry/exit sides, while the DNA around Nuc2 remained relatively wrapped (Figure 3a). In contrast, the NUC_tail_ system maintained a compact architecture, with Nuc1 and Nuc3 consistently closely apposed, which, in turn, was reflected in the small values and narrow Rg distribution as well as small inter-nucleosomal distances (Figure 3b). These observations were further supported by the radius of gyration distribution of the whole system (including all three nucleosomes) (Figure 3c): systems without tails displayed a broad distribution with three peaks that corresponded to various unfolding pathways: single-sided unfolding from Nuc1, unfolding from Nuc3, and simultaneous unfolding from both Nuc1 and Nuc3 (Figure S13). These findings highlight the critical role of histone tails in constraining DNA unwrapping and inter-nucleosome unfolding, preventing the adoption of open conformations, corroborating earlier reports on tail-mediated chromatin condensation ^28–31^.

**Figure 3.**
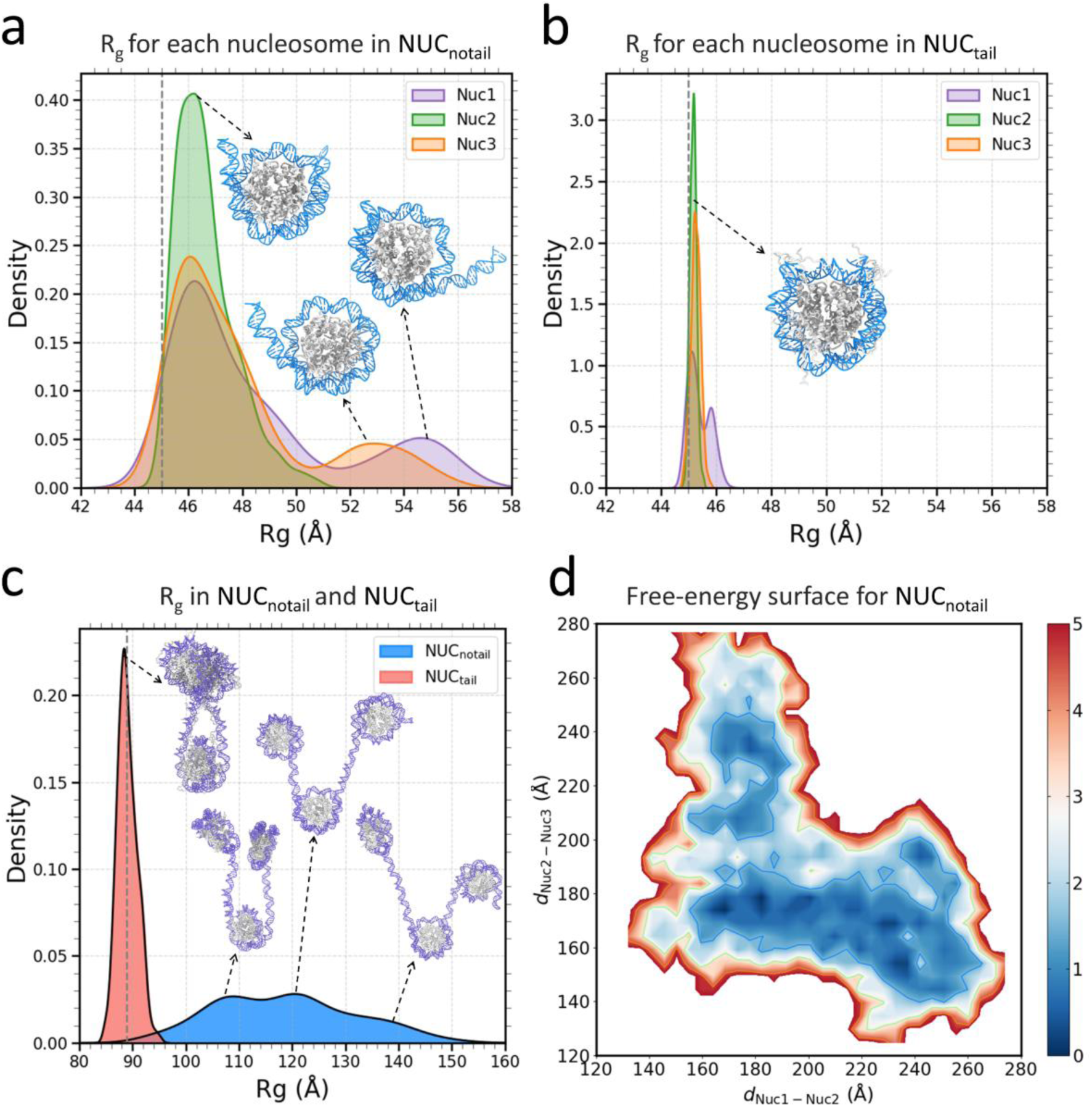
Histone tails mediate tri-nucleosome unfolding and compaction. **a** Radius of gyration (Rg) distributions for each nucleosome (Nuc1: purple, Nuc2: green, Nuc3: red) in the NUC_notail_ system. Grey dashed line shows the Rg values of a single nucleosome in the initial tri-nucleosome system. **b** Same as (a), but for the NUC_tail_ system. **c** Distribution of Rg for all three nucleosomes in NUC_notail_ (blue) and NUC_tail_ (red) tri-nucleosome systems. Grey dashed line shows Rg in the initial tri-nucleosome system. **d** A free-energy surface as a function of center-to-center distances between nucleosome 1 (Nuc1) and nucleosome 2 (Nuc2), and between nucleosome 2 and nucleosome 3 (Nuc3) in NUC_notail_ simulations. Free energies are in kcal/mol.

Next, we tried to interpret our findings by examining the Gibbs free energy landscape as a function of disjointed nucleosomal center-center distances (d_Nuc-Nuc_). In the NUC_notail_ system, the free-energy landscape revealed three wide and dispersed energy minima, aligning with the tri-modal distribution of the radius of gyration of the whole tri-nucleosome system (Figure 3d). Moreover, the energy distribution was wider along the d_Nuc2-Nuc1_ axis compared to the one along the d_Nuc2-Nuc3_ axis. This asymmetry suggests that Nuc1 is more susceptible to unfolding from the middle nucleosome than Nuc3. In contrast, the NUC_tail_ system exhibited a much narrower free-energy basin (Figure S14).

### Molecular Mechanisms of Tail-Tail Mediated Nucleosome Array Compaction

Having established that the tri-nucleosome system with histone tails may capture some aspects of the behavior of compact chromatin, we next sought to elucidate the molecular mechanisms by which histone tails facilitate chromatin compaction at the tri-nucleosome level. Using the NUC_tail_ system, we identified and quantified all inter-nucleosomal histone-histone and histone-DNA interactions. The vast majority of these interactions were mediated by histone tails, specifically by interactions between H2A and H4 tails (forming 23 residue contacts on average) and H3 and H4 tails (12 contacts on average) (Figure S15). Among the disjointed inter-nucleosomal interactions, two types of tail-tail contacts were particularly prevalent between Nuc1 and Nuc3. The first type involved interactions between the H3 tail of Nuc1 and the H4 tail of Nuc3 (Figure 4a). Specifically, salt bridge formation was observed between positively charged residue K9 of the H3 tail and negatively charged residue D24 of the H4 tail. Contact map analyses revealed that this salt bridge interaction was present in approximately 50% of simulation frames (Figure 4b, see Figure S16 for contact maps of different simulation runs). Additional residues involved in H3–H4 tail-tail interactions included T6, R8, and T11 from H3, and L22, Q27, and T30 from H4, with contact frequencies ranging from 10% to 45%.

**Figure 4.**
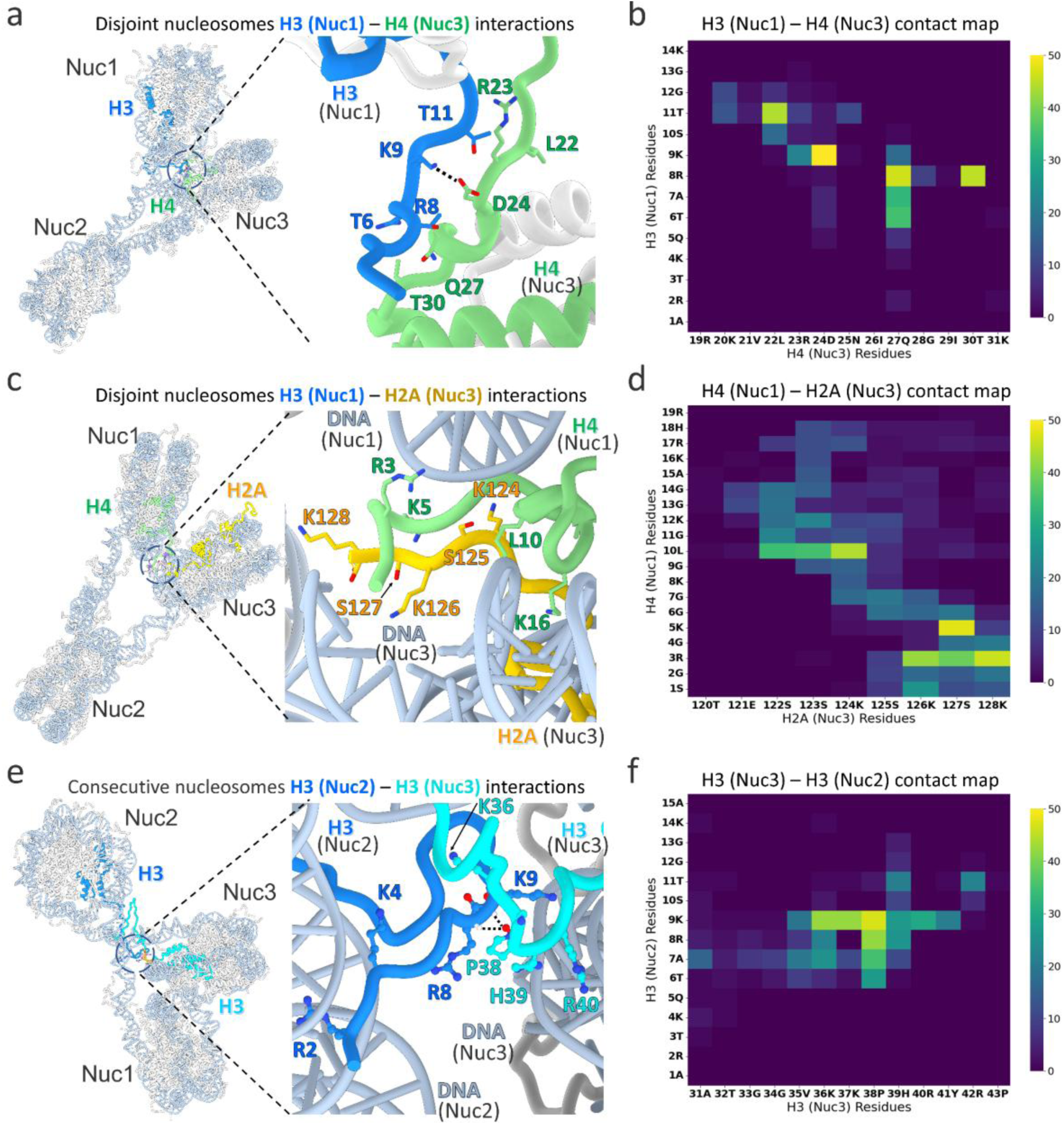
Disjointed and consecutive nucleosomal tail-tail interactions mediate tri-nucleosome compaction. **a** Representative disjointed nucleosomal interactions between the H3 N-terminal tail of Nuc1 and the H4 N-terminal tail of Nuc3, showing key interacting residues. **b, d, f** Contact maps show the percentage of all interactions in MD simulation frames between the corresponding tails in all three independent runs. **c** Disjointed nucleosomal interactions between the H4 N-terminal tail of Nuc1 and the H2A C-terminal tail of Nuc3 via DNA-mediated bridging. **e** Consecutive nucleosome interaction between the H3 N-terminal tails of Nuc2 and Nuc3.

The second type of disjointed nucleosomal interaction was observed between the two DNA gyres of neighboring nucleosomes mediated by histone tails. Specifically, the positively charged residues R3 and K16 from the H4 tail and K124, K126, and K128 from the H2A tail were inserted into the DNA grooves of Nuc1 and Nuc3 respectively, and partially neutralized the DNA negative charge so that DNA gyres of neighboring nucleosomes could come close to each other in space (Figure 4c-d, Figure S17). These tail-mediated dual DNA interactions (a single tail interacting with two DNA regions from different nucleosomes) can promote close nucleosome contacts. We also identified a distinct type of consecutive nucleosomal interaction mediated by the H3 tails from different nucleosomes, which also follows the logic of neutralizing the DNA charge of neighboring nucleosomes and bringing them together (Figure 4e). For instance, the H3 tail of Nuc2 interacted with the linker DNA and the nucleosomal DNA of Nuc3. Key residues included R2, K4, and K9, which were inserted into the minor grooves of the linker and nucleosomal DNA (Figure 4e). Notably, hydrogen bonding was observed between P38 of the H3 tail from Nuc3 and the backbone of R8 and K9 from the H3 tail of Nuc2, with these interactions present in 30% to 45% of simulation frames (Figure 4f, Figure S18). As can be seen, these transient tail–tail interactions were predominantly mediated by lysine residues, whereas arginines were more stably inserted into the DNA minor grooves, as observed in our previous simulations ^32,33^.

### Molecular Mechanisms of Tail-DNA Mediated Nucleosome Array Compaction

In addition to mediating inter-nucleosome interaction through tail-tail contacts, histone tails are known to participate in interactions with DNA that restrict DNA breathing motions and may promote inter-nucleosome interactions ^22,33–38^. Intrinsically disordered histone tails can bind and unbind to specific nucleosomal DNA regions and exist in a dynamic equilibrium, often referred to as fuzzy interactions with DNA ^1,22,32^. To investigate this binding mode further, we mapped all intra- and inter-nucleosomal histone-DNA interactions across the entire 485 bp DNA region of the tri-nucleosome system. Each nucleosome (Nuc1, Nuc2, and Nuc3) in the tri-nucleosome system was assigned its own DNA superhelical location (SHL) positions, including linker DNA, ranging from −SHL8 to +SHL8. Figures 5a and 5b show the mapping of the mean number of H4 tail-DNA and H3 tail-DNA contacts onto the three DNA regions of the tri-nucleosome system (see Figure S18 for H2A and H2B tails). The gray-shaded areas indicate the intra-nucleosomal tail-DNA contacts, while the non-shaded areas represent inter-nucleosomal tail-DNA contacts.

**Figure 5.**
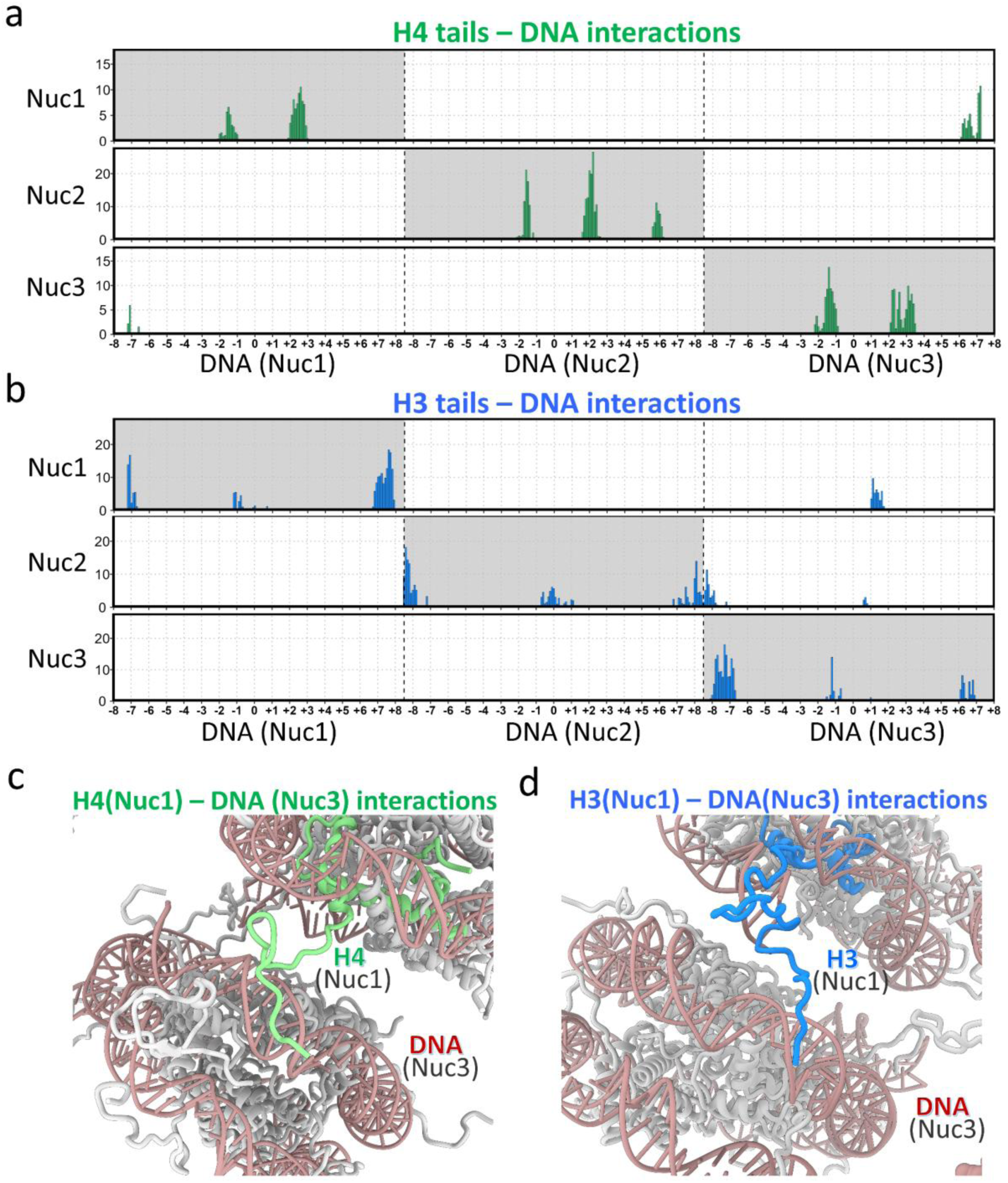
Histone tail - DNA interactions in tri-nucleosome simulations. **a** The mean number of contacts over all MD frames of the three independent runs between the H4 tails from Nuc1, Nuc2, Nuc3 with DNA plotted against the DNA superhelical locations (SHL) in NUC_tail_ nucleosome simulations. Zero corresponds to the dyad position in each nucleosome. **b** The mean number of contacts between the H3 tails from Nuc1, Nuc2, Nuc3 with DNA plotted against DNA SHL in NUC_tail_ nucleosome simulations. **c** A representative conformation of the H4 N-terminal tail from Nuc1 inserting into the minor groove of DNA (SHL+6) from Nuc3 that mediates the disjointed nucleosomal interactions. **d** A representative conformation of the H3 N-terminal tail from Nuc1 interacting with DNA at SHL+1 in Nuc3 that mediates the disjointed nucleosomal interactions. The H3 and H4 tails are highlighted in blue and green, respectively, and the DNA is shown in orange.

First, we showed that the intra-nucleosomal tail-DNA interaction patterns were qualitatively similar across the three nucleosomes (Figure S19). Overall, with a few exceptions, these patterns closely mirrored those found in our earlier mono-nucleosome simulations, which involved a more extensive tail conformational sampling ^22^ that was difficult to achieve in the tri-nucleosome system on this time scale. Because of the nucleosome’s twofold pseudo-symmetry, good enough conformational sampling is expected to yield similar DNA region coverage for each histone copy (negative and positive SHL regions), which we indeed observed for major peaks in the current study. Notably, different histone tails exhibited distinct preferences for different DNA regions. The H2A N-terminal tails were bound primarily to the nucleosomal DNA at superhelical locations (SHL) ± 4, whereas the C-terminal tails of H2A favored SHL ± 7 and the dyad region. The H3 tails, being the longest, engaged multiple DNA regions, including sites near the dyad, and extended up to the next consequent nucleosome via contacts with the linker DNA.

Interestingly, we observed that the H3 and H4 tails from one nucleosome could interact with the DNA region of another nucleosome. For example, H3 and H4 tails from Nuc1 were found to engage with the DNA of Nuc3 by inserting into the DNA minor groove (Figures 5c-d). As for H2A and H2B tails, we observed very few interactions between their respective tails and DNA from other nucleosomes (Figure S18). This indicates that the H2A and H2B tails might play a relatively minor role in compacting the nucleosome arrays through interactions with the DNA of neighboring nucleosomes. However, it should be mentioned that variant nucleosomes containing H2A.Z C-terminal tails have a different sequence and structure, which may extend to the neighboring nucleosomes and form interactions with their acidic patch and mediate higher-order chromatin compaction, as was suggested previously ^39,40^.

## Discussion

Most recent super-resolution microscopy studies showed that nucleosomes are often arranged in discrete and irregular clusters (“clutches”) containing anywhere from a few to a couple of dozen nucleosomes that form a higher-order unit of chromatin organization, particularly in mammalian cells ^41–43^. In this study, using all-atom MD simulations of tri-nucleosomes, which represent structural units of nucleosome clutches, we provide an integrative structural framework that connects atomistic simulations with Cryo-ET map–based chromatin reconstructions. These results establish tri-nucleosome fitting as a new paradigm for understanding native chromatin plasticity at the level of nucleosome clutches and its regulation by histone tails at the atomistic level. Traditionally, chromatin modeling from Cryo-ET/EM maps has relied on fitting mono-nucleosome structures into the local electron density regions. However, such an approach either leaves nucleosomes unconnected or connects them using straight DNA linkers that disregard the geometric and torsional constraints of DNA. These issues are exacerbated in highly condensed chromatin regions, where ambiguous fits and poor DNA resolution complicate the generation of reliable models. Moreover, atomistic-level modeling is limited because histone tail conformations are typically not resolved in mono-nucleosome PDB structures and are therefore omitted in conventional modeling approaches.

Our study demonstrates that fitting tri-nucleosome conformations, sampled from physically realistic MD trajectories, into Cryo-ET maps in some cases may resolve these limitations. The MD conformations not only maintain spatial coherence among three consecutive nucleosomes but also include linker DNA that reflects the intrinsic torsional mechanics and geometric constraints of DNA in chromatin ^44–46^. As a result, tri-nucleosome-based modeling may improve the accuracy and interpretation of nucleosome clutches. This approach also reduces manual intervention during the model construction by embedding disjointed nucleosomal relationships and linker DNA geometry directly into the fitting template, especially valuable in regions where the linker DNA density is missing or partially resolved.

Beyond structural fitting, our tri-nucleosome simulations reveal distinct structural and dynamic features of nucleosome clutches. Histone tails have long been recognized as essential contributors to higher-order chromatin folding, self-association and compaction. It has been shown that even high concentrations of divalent cations, like magnesium, are unable to induce self-association of nucleosomal arrays without histone tails ^47,48^. Indeed, the intrinsically disordered nature of histone tails and their enrichment in positively charged residues enable them to neutralize the negative charge of DNA, thereby facilitating regular chromatin fiber compaction and nucleosome-nucleosome interactions ^48–51^. In addition, the NUC_tail_ simulations revealed that the presence of histone tails was associated with the compaction of tri-nucleosomes, as evidenced by narrow R_g_ distribution and the conformational free-energy landscapes with narrow energy wells, confirmed in different independent runs. To complement previous studies, we provided an atomistic analysis of tail-mediated compaction of nucleosome clutches. It was observed that most tail-tail interactions involved lysines from H2A-H4 and H3-H4 pairs, forming salt bridges and hydrogen bonds, maintained in some cases on microsecond time scale.

Consecutive nucleosomal interactions, particularly those involving H3 tails, further reinforced tri-nucleosome compaction. Previous studies have shown that the H3 tail predominantly stabilizes consecutive nucleosomal interactions ^52,53^ and the absence of histone H3 tails leads to more extended chromatin structures ^29,30,34,54^. Our data revealed that H3 tails can span our linker DNA regions to form extensive networks of tail-tail and tail-DNA interactions. Interestingly, important post-translationally modification sites, like H3K9, were mediating a substantial fraction of these interactions, and, as was shown previously, its methylation may increase the interaction with *i±2* nucleosomes and facilitate the chromatin compaction ^55^.

On the other hand, the NUC_notail_ system, lacking histone tails, exhibited pronounced unfolding that was also characterized by extensive DNA unwrapping, increased disjointed and consecutive inter-nucleosomal distances, and larger effective linker DNA lengths. All these observations confirm previous works ^36,56–58^ and extend them by showing which tail-tail and tail-DNA interactions may mediate inter-nucleosome interactions, their spacing and dynamics. It should be mentioned that the tail-mediated inter-nucleosomal interactions are transient and allow for local dynamical behavior even in the compact state.

Previous *in situ* nucleosome interaction capture experiments in human interphase cells showed that the nucleosomes are mostly engaged in *i±2* interactions (nucleosome *i* with nucleosome *i+2* or nucleosome *i–2*) ^59–62^. The *i±2* interaction patterns are consistent with the two-start zigzag folding of the nucleosome array (in which the nucleosome cores form two twisted columns in there each odd nucleosome (1, 3, 5… etc.) is proximal to the nearest odd nucleosome and each even nucleosome (2, 4, 6… etc.) is proximal to the nearest odd nucleosome) and are especially prominent in repressed heterochromatin. This suggests that the two-start folding is associated with gene repression, consistent with its role in directing heterochromatin-specific interactions with DNA-binding pioneer proteins ^63^. The zigzag chromatin folding has also been observed by *in situ* Cryo-ET at the nuclear periphery ^23,64^. Our tri-nucleosome system, which recapitulates the zigzag interaction motif by interaction between nucleosomes 1 and 3, thus highlights the role of histone tails in bringing the nucleosomes N1 and N3 together. The other feature of the tightly compact zigzag structure is the closely apposed linker DNA exiting and entering nucleosome N2, corresponding to the nucleosome linker “stems” abundant in the repressed chromatin ^65^. Together, these two sites of nucleosome contacts provide the near-atomic details of molecular interactions that can mediate chromatin compaction.

In conclusion, our study introduces a new strategy for integrating atomistic MD simulations with Cryo-ET data to reconstruct chromatin structure at near-atomistic resolution with inherently realistic DNA torsional constraints and inter-nucleosomal relationships. This might help to resolve ambiguities in Cryo-ET maps, particularly in condensed chromatin regions where linker DNA is unresolved, and to capture biologically meaningful nucleosome connectivity. Although the sampling of disordered histone tails in such a large system is limited due to computational complexity, our study is the first to highlight their critical role in modulating tri-nucleosome dynamics and their potential contribution to chromatin compaction at the atomistic scale. These findings provide a mechanistic basis for understanding chromatin plasticity and its functional implications.

## Materials and Methods

### Tri-nucleosome model construction

The tri-nucleosome models used in this study consist of three nucleosome core particles (called “nucleosomes” thereafter) connected by two linker DNA segments (Figure S20). The electron microscopy (EM) structure of the tri-nucleosome (PDB ID: 6L4A) was used as the template for constructing the simulated initial models ^66^. The original DNA sequence from the Cryo-EM structure was replaced with a native DNA sequence generated using Web3DNA ^67^. This native DNA sequence was taken from the *Homo sapiens TP53* gene (GenBank ID: EU877045.1, range: 2486 to 2970) and spans 485 base pairs (bp), with a 147-bp DNA segment wrapping around each nucleosome and two 22-bp DNA linkers connecting nucleosomes. We have constructed two initial tri-nucleosome models: with histone tails (NUC_tail_) and without tails (NUC_notail_). The initial configurations of histone tails (except H3 tails) were taken from the high-resolution X-ray structure of the nucleosome core particle (PDB ID: 1KX5) ^68^. The H3 histone tails were linearly extended symmetrically (with respect to the dyad axis) into the solvent to prevent clashes between nucleosome 1 (Nuc1) and nucleosome 3 (Nuc3). In this study, histone tail regions are defined as follows: H3 N-terminal tail (residues 1-43), H4 N-terminal tail (residues 1-23), H2A N-terminal tail (residues 1-15) and C-terminal tail (residues 119-124), and H2B N-terminal tail (residues 1-29).

### Molecular dynamics simulation protocol

All MD simulations were performed with the package GROMACS version 2022.3 ^69^, using the AMBER 14SB force field for protein parameters and OL15 for nucleic acids ^70,71^, along with the Optimal Point Charge (OPC) water model. This water model has been shown to accurately reproduce bulk water properties and improve the accuracy of simulations involving nucleic acids and intrinsically disordered proteins ^72^. In each simulation, the initial tri-nucleosome model was solvated in a box with a minimum distance of 20 Å between the nucleosome atoms and the box edges. Sodium chloride (NaCl) was added to the system to achieve a concentration of 150 mM.

Energy minimization was performed using the steepest descent algorithm for 10,000 steps. The system was then gradually heated to 300 K over 800 ps, with restraints applied to the atoms, followed by 1 ns of equilibration. Production simulations were conducted in the isobaric-isothermal (NPT) ensemble for up to 3 µs. Temperature was maintained at 300 K using the Bussi thermostat, and pressure was controlled at 1 atm using the Parrinello-Rahman barostat. A cutoff of 10 Å was applied to short-range non-bonded vdW interactions. The Particle Mesh Ewald (PME) method with a cut-off of 10 Å was used to calculate all long-range electrostatic interactions. Periodic boundary conditions were applied, and covalent bonds involving hydrogen atoms were constrained using the LINCS algorithm ^73^, allowing for a 2 fs integration time step. The coordinates of each solute were recorded every 100 ps and a total of 30,000 frames in each simulation run were collected for analysis. For both NUC_notail_ and NUC_tail_ systems, three independent simulation runs were performed.

### Cryo-electron tomographic maps

Cryo-ET maps of chromatin from human K562 cells under three experimental conditions (0 mM Mg²⁺ and 0.75 mM Mg²⁺, *in situ* crosslinked at 0 mM Mg²⁺) were obtained from previous work ^12^. To aid 3D visualization and to improve the resolution of individual nucleosome orientations, we processed the tomograms using deep learning-based regression models ^74^. The chromatin samples, vitrification conditions, and resulting cryotomograms are listed in the Suppl. Table 1 of ^12^ and the raw tilt series are available from Dryad data depository (https://datadryad.org doi:10.5061/dryad.ttdz08m21). Samples analyzed in this work correspond to the tilt series TS21_1 (0 mM MgCl_2_), TS22_1 (0.75 mM MgCl_2_), and TS2_1 (*in situ* crosslinked at 0 mM MgCl_2_).

### Fitting of MD simulation conformations to experimental Cryo-ET maps

To construct three-dimensional (3D) chromatin models at near-atomic resolution, first we fitted the atomic coordinates of the mono-nucleosome structure (PDB: 1KX5) into each denoised Cryo-ET density map (7 maps for chromatin array vitrified at 0 mM Mg^2+^, 13 maps vitrified at 0.75 mM Mg^2+^, 15 maps for *in situ* crosslinked) using the *fitmap* tool in ChimeraX 1.7, which performs rigid-body local optimization ^75^. The voxel size was set to 8.4 Å for the 0 mM Mg²⁺ map (EM magnification 42,000), 5.51 Å for the 0.75 mM Mg²⁺ map (EM magnification 64,000) and 6.65 Å for the crosslinked map (EM magnification 53,000). The *fitmap* tool was configured with a resolution of 16, a cluster angle of 200, and a cluster shift of 24. Initial random placements (1000 in total) were performed of the initial structural model to the 3D density map and then the optimal orientation and position were searched for that maximized the cross-correlation between the model’s density and the experimental map.

In detail, the atomistic model was converted into a density map by representing every atom by a scattering potential approximated by a Gaussian distribution. A global search comprising 1000 random initial placements was performed to explore the full map volume, followed by the local rigid-body optimization to maximize the map–map correlation coefficient. Distinct fitting conformations were identified by clustering the resulting fits using angular and translational thresholds of 200° and 24 Å, respectively. Based on this protocol, we performed the fitting of tri-nucleosome conformations from the MD simulations to native chromatin Cryo-ET maps. Due to the computational cost of the fitting procedure (a large number of conformations, 30,000 frames per simulation × 3 runs × 2 tri-nucleosome systems), we selected 600 MD frames for each NUC_tail_ and NUC_notail_ system, resulting in 600 × 2 × 35 = 42,000 fitting operations. The same *fitmap* parameters were used as in the mono-nucleosome fitting. The MD conformation with the highest goodness-of-fit correlation coefficient value was retained. The correlation coefficient shows the similarity between the experimental Cryo-ET density map and the density map calculated from the atomic model during rigid-body fitting and is defined as:

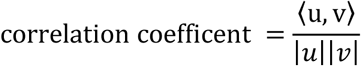

Here, *u* represents the density values of the fit map generated from the atomistic nucleosome model. *v* represents the density values of the reference experimental Cryo-ET map interpolated at the same spatial positions as the grid points. Additionally, the “inside-the-contour” values were calculated, which represent the proportion of atoms from the model that lie within the experimental Cryo-ET map’s contour surface. To calculate the global measures of nucleosome array compaction (disjointed and consecutive inter-nucleosome distances) from experiments, distances between the centers of nucleosomal disks were calculated after fitting these maps with the mono-nucleosomal structural models, as was done in the previous study ^12^.

### Analysis of MD simulations

The MD trajectory frames of each tri-nucleosome system were first aligned by performing a root mean square deviation fitting of the middle nucleosome (Nuc2) in each frame to the corresponding Nuc2 in the energy-minimized tri-nucleosome structure. The fitting was based solely on the Cα atoms of the histone core of Nuc2. For both the NUC_notail_ and NUC_tail_ systems, the initial 200 ns of each simulation trajectory were discarded as for equilibration. The resulting trajectories were visualized using VMD 1.9.3 and analyzed with a set of custom Tcl and Python scripts. Histone-histone and histone-DNA contacts were defined as interactions between any two non-hydrogen atoms within 4.0 Å of each other. The radius of gyration of the DNA segments was calculated using the GROMACS tool gmx_gyrate ^69^. The multi-dimensional free energy landscapes were computed using another GROMACS tool gmx_sham, which estimates the Gibbs free energy by Boltzmann inverting frequency histograms. The nucleosome center-center distances were used as collective variables.

## Acknowledgments

S.L. and A.R.P. were supported by the Department of Pathology and Molecular Medicine, Queen’s University, Canada. A.R.P. is the recipient of a Senior Canada Research Chair in Computational Biology and Biophysics and a Senior Investigator Award from the Ontario Institute of Cancer Research, Canada. A.R.P. and S.L. also acknowledge the support of the Natural Sciences and Engineering Research Council of Canada (NSERC) (No. RGPIN/02972-2021). M.J.A. was supported by a Faculty of Arts and Science infrastructure and a Research Initiation grant. This research is supported by New Frontier in Research Fund Exploration (NFRFE-2021-00880) and Cancer Research Society Operation grants (1056783) to A.R.P. and M.J.A. S.A.G. acknowledges the support of the US National Science Foundation grant 2521597. This study used high-performance computational resources from the Digital Research Alliance of Canada (https://ccdb.alliancecan.ca/).

